# Comparing the microbial communities of natural and supplemental nests of an endangered ecosystem engineer

**DOI:** 10.1101/727966

**Authors:** Megan S. Thoemmes, Michael V. Cove

**Author notes:** Corresponding author: Megan S. Thoemmes, Ph.D., Postdoctoral Scholar. Present address*: Department of Pediatrics, University of California San Diego School of Medicine, Biomedical Research Facility II, 9500 Gilman Drive, La Jolla, CA 92037, USA.

## Abstract

Supplemental nests are often used to restore habitats for a variety of rare and endangered taxa. However, though supplemental nests mimic the function of natural nests, they vary in design and building material. We know from previous research on human homes and other buildings that these differences in architecture can alter the types of microbes to which inhabitants are exposed, and these shifts in microbial interactions can be detrimental for health and well-being. Yet, no one has tested whether microbial communities in supplemental structures are distinct from those found in natural nests. Here we sampled the bacteria from inside supplemental nests of the endangered Key Largo woodrat (*Neotoma floridana smalli*). We then compared the diversity and composition of those bacteria to those collected from natural stick-nests and the forest floor in Key Largo, Florida. In addition, we sampled woodrat bodies to assess the microbiota of nest inhabitants. We observed distinct bacterial communities in Key Largo woodrat nests, relative to the forest environment; however, we could not differentiate between the microbial communities collected from supplemental and natural nests. Furthermore, when we considered genera known to contain bacterial pathogens of wild rodents, supplemental and natural nests exhibited similarly low relative abundances of these taxa. Where we expected to see an accumulation of pathogens, we instead observed high relative abundances of bacteria from antimicrobial-producing groups (i.e., Pseudonocardiaceae and Streptomycetaceae). The microbial biota of Key Largo woodrat individuals resembled those of their nests, with low relative abundances of potentially pathogenic bacteria and high abundances of antimicrobial-producing groups. Our results suggest that, although there is some microbial interaction between nests and nest inhabitants, there are no detectable differences in the types of bacteria to which Key Largo woodrats are exposed in supplemental and natural nest structures.

## Introduction

Supplemental nests are often used in the conservation and management of threatened and endangered species to increase and restore available nesting habitat and to provide protection from predators, competitors, and the environment (Newton, 1994; Spring et al., 2001; Libois et al., 2012). Though supplemental nests model the function of their natural counterparts, they are commonly built from manufactured materials, with relatively little attention to mimicking the details of natural nest design. However, we know from the study of other human-built structures (e.g., homes and office buildings) that building material and architectural design strongly influence the diversity and types of microbes found on interior surfaces. Contemporary houses have far fewer environmental microbes than do more open traditional homes (Ruiz-Calderon et al., 2016, Thoemmes et al., *In Prep*), which have fewer environmental microbes than do chimpanzee nests (Thoemmes et al., 2018). While the absence of some bacteria in our daily lives is beneficial, the absence of others is associated with negative health outcomes. For example, a decrease in the abundance of soil bacteria on the skin is directly linked to an increase in the prevalence of atopic sensitization and autoimmune disorders in humans (Fyhrquist et al., 2014; Ruokolainen et al., 2015). Additionally, as indoor microbial diversity decreases there is a subsequent increase in the abundance of bodily microbes, such as those from feces and skin (Dunn et al., 2013; Lax et al., 2014), as well as pathogenic bacteria in both homes and hospitals (Kembel et al., 2014). Despite this, no one has ever studied how human-built supplemental nests alter the microbial communities to which other species are exposed. A loss in microbial diversity or the accumulation of pathogens in supplemental nests could have detrimental effects, particularly for species at high risk of extinction, such as the Key Largo woodrat (*Neotoma floridana smalli*).

Key Largo woodrats are a federally endangered subspecies endemic to north Key Largo, FL (US Department of the Interior., 1984). These woodrats are ecosystem engineers and modify their environment by building substantial natural stick-nests that are maintained across generations by layering sticks and debris at the bases of trees, in fallen tree throws, or in solution holes (Cove and Maurer, 2019). Once estimated to number fewer than 100 individuals (McCleery et al., 2005), Key Largo woodrats have benefitted greatly from adaptive management, including nest supplementation, due to limited natural nesting substrate from historical land alterations (Cove et al., 2017). In fact, there have now been more than 2000 supplemental Key Largo woodrat nests built in the last remaining upland hammock habitat of the Crocodile Lake National Wildlife Refuge and Dagny Johnson Botanical State Park (Cove et al., 2017). Supplemental nests differ from natural nests in that they are constructed from large culvert pipes covered with rocks or chunks of fossilized coral. On the exterior, natural and supplemental nests are maintained in similar ways through stick-stacking behavior (Cove et al., 2017), but it is on the interior of nests where differences become more apparent. Supplemental nests are enclosed with comparatively little air flow and moisture penetration (Barth, 2014). This could limit the dispersal of environmental species into nests and alter microclimate conditions, affecting the overall diversity and succession of microbial communities.

Here we examine the diversity and composition of bacteria in natural and supplemental Key Largo woodrat nests to assess whether there are differences in nest microbiomes associated with nest supplementation. Based on what we know from human homes, we might expect supplemental nests to have less bacterial diversity. In addition, as humans have sealed ourselves off from the outdoor environment, we have observed a shift in the relative abundance of microbial taxa (Adams et al., 2014; Miletto and Lindow, 2015; Barberán et al., 2015; Thoemmes et al., 2018), where we see an increase in the accumulation of body microbes and pathogenic bacteria (e.g., *Staphylococcus aureus*; Gandara et al., 2006). Since supplemental nests are composed from materials that could restrict the colonization of environmental microbes, we might expect there to be a difference in which bacterial taxa are most abundant when compared to natural nests. Finally, as we know there is an interaction between the microbiota of the body and the built environment (Hospodsky et al., 2012; Dunn et al., 2013; Meadow et al., 2014; Lax et al., 2014; Gibbons et al., 2015), the composition of bacterial communities in nests might be influenced by the nest inhabitants themselves. Therefore, we characterize the bacteria found on the bodies of Key Largo woodrats.

## Methods

The Crocodile Lake National Wildlife Refuge is located in North Key Largo, FL, USA. Though the majority of this refuge is composed of mangroves and coastal wetlands, it is part of the last remaining large tract of tropical hardwood hammock forest. When combined with the Dagny Johnson Key Largo Hammock Botanical State Park, this forest type covers less than 1000 ha (Frank et al., 1997; US Fish and Wildlife Service, 1999) and is home to a variety of endemic and endangered species, including Key Largo woodrats, Key Largo cotton mice (*Peromyscus gossypinus allapaticola*), and Stock Island tree snails (*Orthalicus reses*). Our study focused on the Key Largo woodrat, in which we conducted all research in December 2017.

Previously located natural nests and all supplemental nests are individually marked as part of a long-term monitoring project. From these, we visited 10 natural and 10 supplemental nests (n = 20) located in the area of the refuge that exhibits the highest population densities of Key Largo woodrats (Cove et al., *In review*). We determined nest occupancy based on visual surveys of active stick-stacking behavior (Balcom and Yahner, 1996; Cove et al., 2017) or additional evidence of use (such as with camera trap surveys or Sherman live-traps) and swabbed each nest with dual-tipped sterile BBL™ CultureSwabs™. To standardize the distance into each nest, as well as to avoid contamination of sample swabs on exterior building material, we inserted a PVC pipe (approximately 0.5 m in length) into each nest prior to sample collection. We targeted areas that appeared to be used most frequently by the inhabitant(s) and then thread each swab through the pipe at the sample site. To determine how bacterial communities in nests vary, relative to the surrounding forest environment, we then swabbed the forest floor ~0.5-0.75 m from the exterior of natural nests (n = 10), in an area that did not appear to be trafficked by humans or wildlife. We swabbed all nest and forest floor environments for approximately 15 seconds.

In addition, as we know there is an interaction between mammal bodies and the structures they inhabit (Pakarinen et al., 2008; Hospodsky et al., 2012; Hanski et al., 2012; Dunn et al., 2013), we characterized the bacteria that live on the bodies of the Key Largo woodrats. As part of the long-term monitoring efforts, we captured individual woodrats near sampled nests with Sherman live-traps and swabbed their flank and ventral side (n = 10) for approximately 15 seconds. To prevent repeated sampling of individuals, we verified identity with double-marked monel #1005 ear (National Band and Tag Company, Newport, KY) and subcutaneous PIT (Biomark, Boise, ID) tags.

### Molecular Methods and Analyses

We performed DNA extractions with a DNeasy PowerSoil Kit (Qiagen, product #12888-100), with modifications described in Fierer et al. (2008), and included two sterile swabs in our extraction process to identify potential contaminants. PCR reactions were performed by the Fierer lab (University of Boulder, Colorado) in triplicate and all amplicons were pooled in equimolar concentrations prior to sequencing on the Illumina MiSeq platform, where we targeted the V4-V5 region of the 16S rRNA bacterial gene (Flores et al., 2012).

We processed all data using default parameters in the QIIME v1.9.1 pipeline (Caporaso et al., 2010) and picked operational taxonomic units (OTUs) with UCLUST at a 97% similarity (DeSantis et al., 2006; Edgar, 2010). Taxonomy was assigned with the RDP Classifier (Wang et al., 2007), and we removed all resulting OTU sequences from downstream analyses that were classified as mitochondria, chloroplast, and unassigned, as well as OTUs that were amplified from control samples (i.e., those from blank swabs and blank reagent wells). To quality filter our data, we used the Bokulich threshold, removing all OTUs that represented less than 0.005% of total read abundance (Bokulich et al., 2013; Nguyen et al., 2015). We then analyzed all data in the R environment with the mctoolsr, vegan, PMCMRplus, and FSA packages (Oksanen et al., 2013; R Core Team, 2015; Leff, 2016; Pohlert and Pohlert, 2018; Ogle, 2018).

We compared differences in OTU richness and Shannon diversity index between nest and forest floor samples with Kruskal-Wallis, using the Dunnett’s test and Benjamini-Hochberg method for multiple comparisons (n = 30; Dunnett, 1955; Benjamini and Hochberg, 1995). We then quantified differences in the composition of bacterial communities among samples with Bray-Curtis dissimilarity, weighted by OTU abundance (Bray and Curtis, 1957). We considered differences in bacterial community composition between natural and supplemental nests with a permutational multivariate analysis of variance (PERMANOVA), where we compared differences between nest type (i.e., natural and supplemental nests) and natural nest and forest samples separately.

We then tested for the presence of bacteria from genera known to contain zoonotic bacterial pathogens of wild rodents, as characterized previously by Razzauti et al. (2015). Though there is likely to be some variation in the pathogens found on Key Largo woodrats compared to other rodent species, these taxa encompass 45 bacterial genera, including those of well-known rodent-associated pathogenic groups (e.g., *Bartonella*, *Rickettsia*, *Borrelia*, *Neoehrlichia*, *Anaplasma*, and *Yersinia pestis*). Therefore, though we may have missed fine-scale interactions (e.g., previously undescribed pathogens or species-specific associations), we believe this representative dataset has captured any potential generalized patterns in the accumulation of pathogenic bacteria in Key Largo woodrat nests. We then calculated the percent relative abundance of all pathogenic genera and compared differences between nest type and forest floor samples with Kruskal-Wallis tests.

To consider the bacteria found on Key Largo woodrat bodies, we characterized the overall percent relative abundance of all bacterial taxa found in nests (n = 20) and on individuals (n = 10). In addition, we compared the accumulation of bacterial taxa that contain pathogenic lineages in nests to those found on individuals. Finally, we compared differences in community composition between nests and bodies with Bray-Curtis dissimilarity (Bray and Curtis, 1957) and visualized community data with non-metric multidimensional scaling (NMDS) ordination plots. We quantified differences with PERMANOVA, using an FDR correction for multiple comparisons.

## Results

We observed a total of 1952 unique OTUs across all natural Key Largo woodrat nests, with an average of 776 OTUs represented per individual nest. This was no different from the OTU richness observed from supplemental nests (average of 919 OTUs per individual nest; *P* = 0.41) or from the forest floor (average of 841 OTUs per sample location; *P* = 0.15; Figure 1). We found similar results when we compared the Shannon diversity index, with no difference between natural and supplemental nests (*P* = 0.56) or between natural nests and the forest floor outside of each nest (*P* = 0.11). We also observed no differentiation in the composition of bacterial community membership between natural and supplemental nests (PERMANOVA: *P* = 0.58; Figure 2); however, we did detect differences in bacterial communities in Key Largo woodrat nests overall compared to the surrounding forest environment (PERMANOVA: *P* < 0.001; Figure 2).

**Figure 1.**
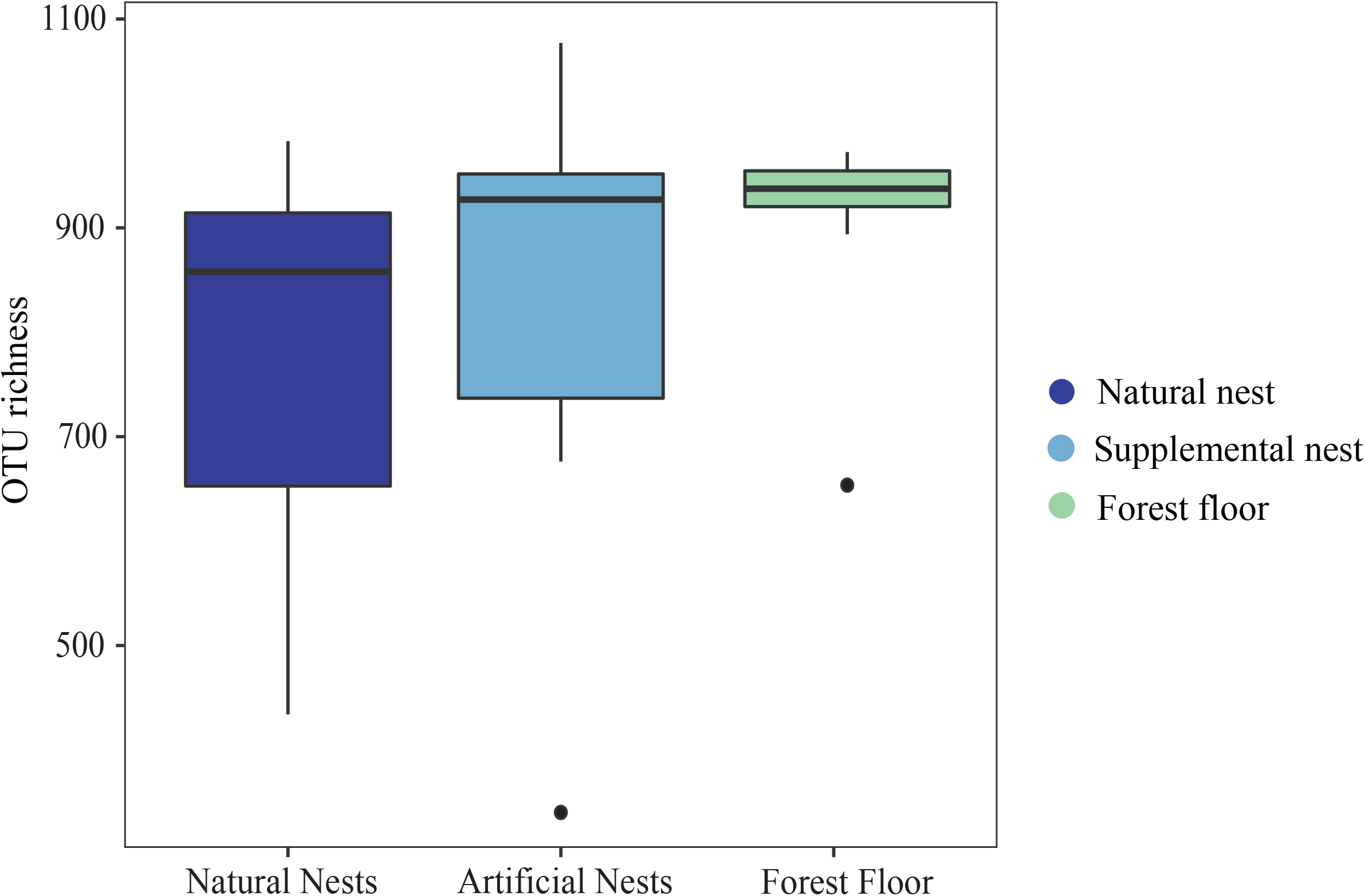
OTU richness of bacteria collected from natural nests, supplemental nests, and the forest floor outside of each natural nest of the Key Largo woodrat. There were no observed differences in OTU richness between sample type (natural and supplemental nests: *P* = 0.41; nests and forest floor: *P* = 0.15).

**Figure 2.**
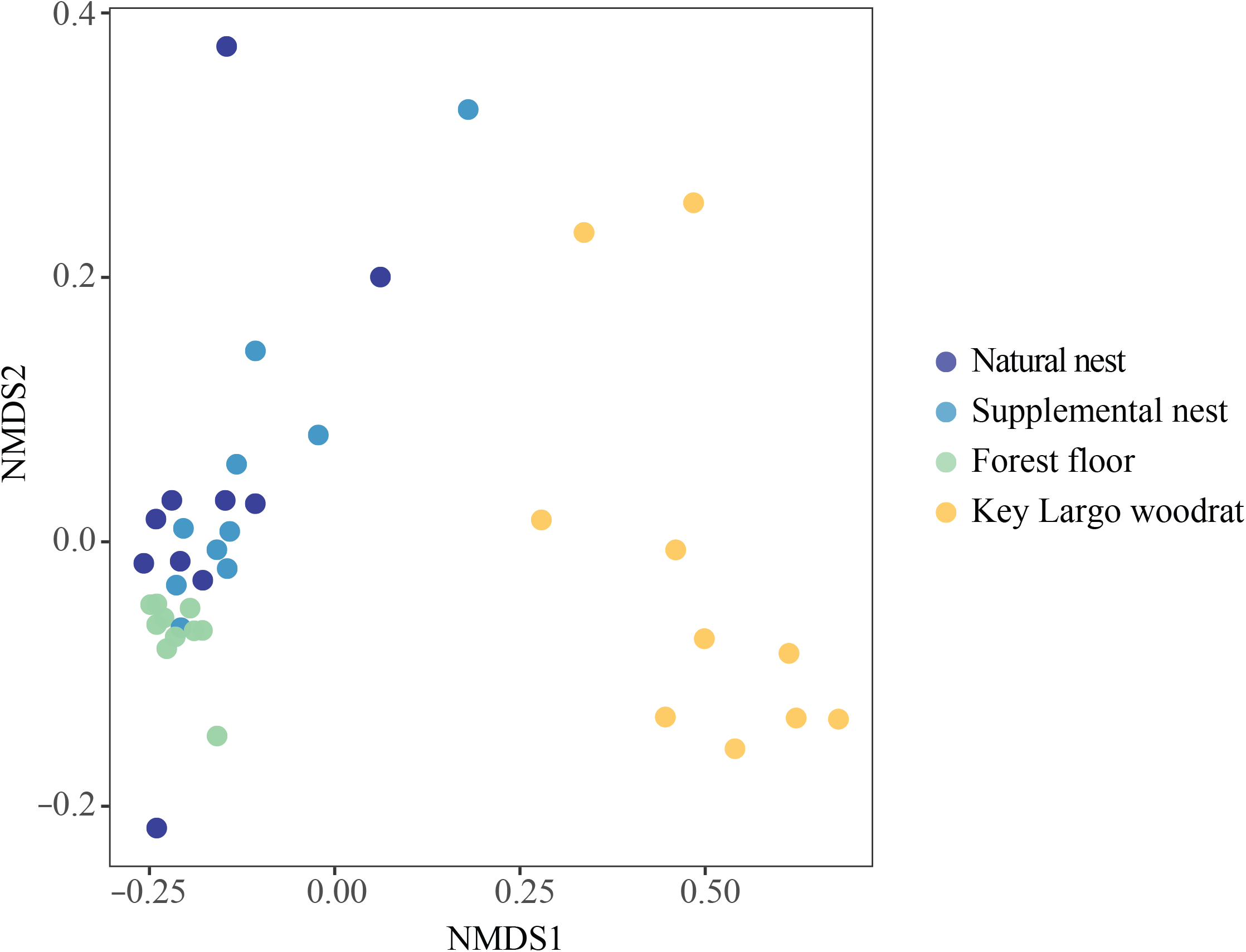
Non-metric multidimensional scaling (NMDS) ordination plot representing bacterial communities collected from natural and supplemental Key Largo woodrats nests, the forest floor outside each natural nest, and Key Largo woodrat bodies. There were distinct bacterial communities in Key Largo woodrat nests compared to the forest floor (*P* < 0.001), but there was no difference between natural and supplemental nests (*P* = 0.58). Key Largo woodrat microbiomes were different from the composition of bacteria found in their nests (*P* < 0.001), and individuals exhibited a greater amount of variation in bacterial communities compared to their nest and forest environments.

Differences among bacterial communities in Key Largo woodrat nests (relative to the forest floor) were not driven by an accumulation of bacteria from pathogenic lineages. The relative abundances of these taxa were only slightly higher in nests than from the forest floor (percent of bacterial sequence reads: natural = 2.6%, supplemental = 3.1%, forest = 1.6%; χ^2^ = 7.93, *P* = 0.005), and there were no more of these pathogenic groups in supplemental nests than in natural nests (χ^2^ =1.801, *P* = 0.18). Of the 45 genera tested that contain rodent-associated pathogens, we detected only 10 in nests. These included *Bacillus*, *Burkholderia*, *Clostridium*, *Corynebacterium*, *Enterococcus*, *Micrococcus*, *Mycobacterium*, *Nocardia*, *Rhodococcus*, and *Vibrio*, eight of which were shared between natural and supplemental nests (Table 1). Furthermore, we found all recovered taxa at relatively low abundances (read numbers), with *Mycobacterium* accounting for the greatest relative abundance at 1.5% of total bacterial sequence reads in natural nest samples and 1.1% of bacterial sequence reads from supplemental nest samples (Table 1). *Mycobacterium* was followed by *Burkholderia* (0.4% of bacterial sequences) in natural nests and *Nocardia* in supplemental nests (0.8% of bacteria sequences), of which only *Nocardia* was more abundant in nests than from forest floor samples (*Burkholderia*: χ^2^ = 0.041, *P* = 0.84; *Nocardia*: χ^2^ = 9.401, *P* = 0.002; Table 1).

**Table 1.** Bacterial genera detected in our samples that contain known rodent-associated pathogens. All values represent the average percent relative abundance of total bacterial sequence reads per sample type. Taxa with the greatest relative abundance are marked with an asterisk (*).

| **Genera** | **Natural** | **Supplemental** | **Forest** | **KL woodrat** |
| --- | --- | --- | --- | --- |
| *Bacillus* | 0.3 | 0.3 | 0.4 | 0.2 |
| *Burkholderia* | 0.4 | 0.1 | 0.07 | 0.04 |
| *Clostridium* | 0 | 0.002 | 0 | 0.1 |
| *Corynebacterium* | 0.004 | 0.1 | 0.002 | 0.6* |
| *Enterococcus* | 0.002 | 0 | 0.002 | 0.2 |
| *Eubacterium* | 0 | 0 | 0.002 | 0 |
| *Haemophilus* | 0 | 0 | 0 | 0.07 |
| *Mannheimia* | 0 | 0 | 0 | 0.03 |
| *Micrococcus* | 0.01 | 0.2 | 0.006 | 0.05 |
| *Moraxella* | 0 | 0 | 0 | 0.002 |
| *Mycobacterium* | 1.49* | 1.12* | 1.46* | 0.4 |
| *Mycoplasma* | 0 | 0 | 0 | 0.002 |
| *Nocardia* | 0.3 | 0.8 | 0.02 | 0.1 |
| *Rhodococcus* | 0.02 | 0.1 | 0.014 | 0.07 |
| *Stenotrophomonas* | 0 | 0 | 0 | 0.01 |
| *Streptococcus* | 0 | 0 | 0.004 | 0.5 |
| *Treponema* | 0 | 0 | 0 | 0.1 |
| *Vibrio* | 0.04 | 0.01 | 0 | 0 |

However, while we saw little accumulation of pathogenic taxa, we saw a high relative abundance of Pseudonocardiaceae and Streptomycetaceae in Key Largo woodrat nests. Both of these families are important antimicrobial-producing groups (Platas et al., 1998; Kämpfer et al., 2014) and include the bacteria that produce many of our common commercial antibiotics, such as erythromycin and vancomycin (a well-known treatment for Methicillin-resistant *Staphylococcus aureus* infections; Sakoulas et al., 2004; Jafari et al., 2014; Kämpfer et al., 2014). Relative to other taxa, Pseudonocardiaceae and Streptomycetaceae were the most abundant bacteria in both natural and supplemental nests, accounting for 12% of all sequence reads from natural nests and 15% of all sequence reads from supplemental nests. And though these taxa are found in diverse environments and are abundant in soils globally, they were notably more abundant in nests compared to the forest floor (only 4% of sequence reads from forest samples; χ^2^ = 9.486, *P* = 0.002).

Due to the accumulation of Pseudonocardiaceae bacteria in nests, we also tested for its presence on the Key Largo woodrat. Just as with their nests, Pseudonocardiaceae was abundant on Key Largo woodrats, where it was the third most abundant bacterial family (7% of bacterial sequences). When we considered the accumulation of pathogenic taxa on individuals, we observed 16 of the 45 genera considered (Table 1), where *Corynebacterium* was the most abundant of these genera (0.6% of total bacterial sequences). *Mycobacterium* was present on woodrats (as well as in nests) and includes noxious pathogens, such as those known to cause tuberculosis and leprosy in mammals (Hansen, 1874; Koch, 1884; Gordon and Parish, 2018). However, they were found at very low abundances (0.4% of bacterial sequences). Overall, the Key Largo woodrat had approximately the same relative abundance of pathogenic taxa (2.4% of bacterial sequences) as was detected in their natural nests (2.9% of bacterial sequences; *P* = 0.415). Additionally, though Key Largo woodrats resembled their nests, in relation to the relative accumulation of pathogenic taxa and antimicrobial-producing bacteria, woodrats were distinctly different from their nests (PERMANOVA: *P* < 0.001), even more so than were nests from the forest environment (PERMANOVA: *P* < 0.001; Figure 2).

## Discussion

Overall, we found bacterial communities in Key Largo woodrat nests to be distinct from the forest environment, but we observed no differences between natural and supplemental nests, based on the diversity or composition of those bacteria. There was little accumulation of bacteria from pathogenic lineages in nests or on Key Largo woodrat individuals. Instead, we found a high abundance of bacteria from antimicrobial-producing groups.

We expected supplemental nests to be similar to other structures built by humans (e.g., contemporary homes and office buildings), in that, they might be expected to have a less diverse, unique assemblage of bacteria compared to natural nests (Dunn et al., 2013; Lax et al., 2014). If supplemental nests alter bacterial species interactions, this could have detrimental effects on woodrat health. The exposure to a greater diversity of microbes increases immune response and the ability to fight off infectious disease in rodents (Beura et al., 2016). However, we found no evidence of such an effect. Relative to the forest floor, nests had similar bacterial diversity, regardless of whether they were natural or supplemental (Figure 1), and we could not discern difference between nest type, based on which bacterial taxa were present (Figure 2). This suggests that, likely through some combination of nest design or pattern of use, supplemental nests maintain a bacterial community that is no different from their natural counterparts. One explanation might be that the culvert pipes used in supplemental nest construction have open ends. These openings could act in a similar way to the gaps in natural nests or to open windows in human homes (Kembel et al., 2012; Barberán et al., 2015).

We also found very little accumulation of bacteria from pathogenic lineages in natural and supplemental nests (Table 1). With the exception of *Mycobacterium*, a genus that contains the bacteria that cause tuberculosis and leprosy (Hansen, 1874; Koch, 1884; Gordon and Parish, 2018; but which also includes a high diversity of harmless species), the most common pathogenic taxa were found at very low abundances (*Burkholderia* and *Nocardia*; less than 1% of total bacteria; Table 1). *Burkholderia* and *Nocardia* are known to contain primarily non-pathogenic species, and further still, pathogens in *Nocardia* are frequently classified as opportunistic. High bacterial diversity and a high relative abundance of Pseudonocardiaceae and Streptomycetaceae in nests are likely contributing factors, as we know that higher bacterial diversity and the application of antimicrobials in human buildings are associated with a decrease in the relative abundance of pathogens (Lax et al., 2014; Ruokolainen et al., 2017). On the other hand, the application of antimicrobials in homes has favored antimicrobial-resistant strains (Hartmann et al., 2016), and therefore, we may expect to find antibiotic-resistant bacteria in Key Largo woodrat nests, particularly since they are typically used for several generations and persist over long periods of time (Rainey, 1956). However, as our molecular methods are not reliable at species level of identification (Martínez-Porchas et al., 2016; Edgar, 2018), it would be imprudent for us to make strong conclusions about the presence of individual species or strains within the scope of this study. Due to this, we cannot directly attribute the bacteria in Key Largo woodrat nests to pathogenesis but rather use this data as a proxy for understanding the broad-scale patterns of pathogen accumulation.

Since we know that the species found in human homes are influenced by the building’s occupants (Barberán et al., 2015), we characterized the bacteria from Key Largo woodrat bodies. Of the pathogenic taxa considered, Key Largo woodrats had the highest relative abundance of *Corynebacterium* (Table 1). Pathogens in *Corynebacterium* can cause disease, such as diphtheria and endocarditis, but this genus contains primarily non-pathogenic species and is a common associate of mammal skin (Loeffler, 1884; et Yersin, 1888; Almklov and Hansen, 1950; Pike, 1951). The other pathogenic genera largely contained opportunistic rodent-associated pathogens (Table 1). The low detection of pathogens in nests and on bodies may be due, in part, to the high abundances of Pseudonocardiaceae and Streptomycetaceae. It is also notable that the differentiation and variation among woodrat microbiomes was much greater than what we observed between nest and forest samples (Figure 2). One potential explanation could be the landfall of Hurricane Irma in September 2017 (3 months prior to sample collection). Catastrophic weather events can homogenize biological communities (Savage et al., 2018), and therefore, the hurricane could account for the similarity between and among nest and forest microbiomes.

One of our more unusual findings was the prevalence of bacteria from the Pseudonocardiaceae and Streptomycetaceae families in Key Largo woodrat nests. Associations between animals and antimicrobial-producing bacteria have been described in social insects (e.g., from ants and wasps; Currie et al., 1999; Cafaro and Currie, 2005; Madden et al., 2013). However, to our knowledge, such a relationship has never been observed from mammals. Based on our study design, we cannot ascribe a causative relationship between the Key Largo woodrats and the presence of these bacteria. However, due to their high abundance and ubiquity among natural and supplemental nests, we propose the possibility that the bodies and/or behaviors of Key Largo woodrats promote the accumulation of these beneficial bacteria.

Unlike other human-built structures, such as homes and hospitals, supplemental nests for the Key Largo woodrat do not appear to alter microbial species interactions in the ways we would predict to be detrimental for woodrat health. However, due to variation in the types of nests and nest boxes used in supplementation studies for threatened and endangered species conservation, we recommend that more research is required prior to the extrapolation of results to the nests constructed for other species of concern.

## Acknowledgments

This research was supported by the Brevard Zoo’s Quarters for Conservation Fund, The Florida Keys Wildlife Society, USFWS, the Florida Keys National Wildlife Refuge Complex, and NC Cooperative Fish and Wildlife Research Unit. Special thanks to J. Dixon, S. Sneckenberger, M. Jee, R. DeGayner, and C. DeGayner for their efforts and continued support building supplemental nests and conducting visual, live-trap, and camera trap surveys.

## Data Accessibility

Microbial data from this project were deposited in the Mendeley Data repository and made publicly available at http://dx.doi.org/10.17632/wsys8nt5h6.1

## Ethics Statement

This research was conducted and all data were collected in accordance with federal, state, and institutional animal ethics guidelines and approved under the following permits: US Fish and Wildlife Service Permit [TE 697819-4]; Florida Department of Environmental Protection [#01171715]; and the North Carolina State University Institutional Animal Care and Use Committee (IACUC) [#13-003-O].

## References

Adams RI, Miletto M, Lindow SE, Taylor JW, Bruns TD. 2014. Airborne bacterial communities in residences: similarities and differences with fungi. PLoS One, 9:e91283.

Almklov JR, Hansen AE. 1950. Successful treatment of *C. diphtheriae* subacute bacterial endocarditis with penicillin and streptomycin. Pediatrics, 5:437–442.

Balcom BJ, Yahner RH. 1996. Microhabitat and landscape characteristics associated with the threatened Allegheny woodrat. Conserv Biol, 10:515–525.

Barberán A, Dunn RR, Reich BJ, Pacifici K, Laber EB, Menninger HL, Morton JM, Henley JB, Leff JW, Miller SL. 2015. The ecology of microscopic life in household dust. Proc R Soc B, 282:20151139.

Barberán A, Ladau J, Leff JW, Pollard KS, Menninger HL, Dunn RR, Fierer N. 2015. Continental-scale distributions of dust-associated bacteria and fungi. PNAS, 201420815.

Barth LJ. 2014. Habitat use of the Key Largo woodrat (*Neotoma floridana smalli*). MS Thesis. Florida International University, FL, USA.

Benjamini Y, Hochberg Y. 1995. Controlling the false discovery rate: a practical and powerful approach to multiple testing. J R Stat Soc Series B Stat Methodol, 289–300.

Beura LK, Hamilton SE, Bi K, Schenkel JM, Odumade OA, Casey KA, Thompson EA, Fraser KA, Rosato PC, Filali-Mouhim A. 2016. Normalizing the environment recapitulates adult human immune traits in laboratory mice. Nature, 532:512.

Bokulich NA, Subramanian S, Faith JJ, Gevers D, Gordon JI, Knight R, Mills DA, Caporaso JG. 2013. Quality-filtering vastly improves diversity estimates from Illumina amplicon sequencing. Nat Methods, 10:57.

Bray JR, Curtis JT. 1957. An ordination of the upland forest communities of southern Wisconsin. Ecol Monograph, 27:325–349.

Cafaro MJ, Currie CR. 2005. Phylogenetic analysis of mutualistic filamentous bacteria associated with fungus-growing ants. Can J Microbiol, 51:441–446.

Caporaso JG, Kuczynski J, Stombaugh J, Bittinger K, Bushman FD, Costello EK, Fierer N, Pena AG, Goodrich JK, Gordon JI. 2010. QIIME allows analysis of high-throughput community sequencing data. Nat Methods, 7:335.

Cove MV, Maurer AS. 2019. Home decorating with an endangered ecosystem engineer. Front Ecol Environ, 17: 231.

Cove MV, Simons TR, Gardner B, Maurer AS, O’Connell AF. 2017. Evaluating nest supplementation as a recovery strategy for the endangered rodents of the Florida Keys. Restor Ecol, 25:253–260.

Cove MV, Simons TR, Gardner B, O’Connell AF. *In revision*. Towards recovery of an endangered island endemic: numerical and behavioral responses of Key Largo woodrats to exotic predator removal. Biol Conserv.

Currie CR, Scott JA, Summerbell RC, Malloch D. 1999. Fungus-growing ants use antibiotic-producing bacteria to control garden parasites. Nature, 398:701.

DeSantis TZ, Hugenholtz P, Larsen N, Rojas M, Brodie EL, Keller K, Huber T, Dalevi D, Hu P, Andersen GL. 2006. Greengenes, a chimera-checked 16S rRNA gene database and workbench compatible with ARB. Appl Environ Microbiol, 72:5069–5072.

Dunn RR, Fierer N, Henley JB, Leff JW, Menninger HL. 2013. Home life: factors structuring the bacterial diversity found within and between homes. PloS One, 8:e64133.

Dunnett CW. 1955. A multiple comparison procedure for comparing several treatments with a control. J Am Stat Assoc, 50:1096–1121.

Edgar RC. 2018. Updating the 97% identity threshold for 16S ribosomal RNA OTUs. Bioinformatics, 1:5.

Edgar RC. 2010. Search and clustering orders of magnitude faster than BLAST. Bioinformatics, 26:2460–2461.

et Yersin R. 1888. Contribution à l’Étude de la Diphtérie. Annales de l’Institut Pasteur.Décbr.

Fierer N, Hamady M, Lauber CL, Knight R. 2008. The influence of sex, handedness, and washing on the diversity of hand surface bacteria. PNAS, 105:17994–17999.

Flores GE, Henley JB, Fierer N. 2012. A direct PCR approach to accelerate analyses of human-associated microbial communities. PloS One, 7:e44563.

Frank P, Percival F, Keith B. 1997. A status survey for Key Largo woodrat (*Neotoma floridana smalli*) and Key Largo cotton mouse (*Peromyscus gossypinus allapaticola*) on North Key Largo, Monroe County, Florida. Unpublished report to the US Fish and Wildlife Service, Jacksonville, Florida, 1–21.

Fyhrquist N, Ruokolainen L, Suomalainen A, Lehtimäki S, Veckman V, Vendelin J, Karisola P, Lehto M, Savinko T, Jarva H. 2014. *Acinetobacter* species in the skin microbiota protect against allergic sensitization and inflammation. J Allergy Clin Immunol, 134:1309. e11.

Gandara A, Mota LC, Flores C, Perez HR, Green CF, Gibbs SG. 2006. Isolation of *Staphylococcus aureus* and antibiotic-resistant *Staphylococcus aureus* from residential indoor bioaerosols. Environ Health Perspect, 114:1859.

Gibbons SM, Schwartz T, Fouquier J, Mitchell M, Sangwan N, Gilbert JA, Kelley ST. 2015. Ecological succession and viability of human-associated microbiota on restroom surfaces. Appl Environ Microbiol, 81:765–773.

Gordon SV, Parish T. 2018. Microbe profile: *Mycobacterium tuberculosis*: humanity’s deadly microbial foe. Microbiology (Reading, England), 164:437–439.

Hansen GA. 1874. Undersøgelser angående spedalskhedens årsager. Norsk Magazin for Laegeviteuskapen, 4:1–88.

Hanski I, von Hertzen L, Fyhrquist N, Koskinen K, Torppa K, Laatikainen T, Karisola P, Auvinen P, Paulin L, Mäkelä MJ. 2012. Environmental biodiversity, human microbiota, and allergy are interrelated. PNAS, 109:8334–8339.

Hartmann EM, Hickey R, Hsu T, Betancourt Román CM, Chen J, Schwager R, Kline J, Brown GZ, Halden RU, Huttenhower C. 2016. Antimicrobial chemicals are associated with elevated antibiotic resistance genes in the indoor dust microbiome. Environ Sci Technol, 50:9807–9815.

Hospodsky D, Qian J, Nazaroff WW, Yamamoto N, Bibby K, Rismani-Yazdi H, Peccia J. 2012. Human occupancy as a source of indoor airborne bacteria. PloS One, 7:e34867.

Jafari N, Behroozi R, Farajzadeh D, Farsi M, Akbari-Noghabi K. 2014. Antibacterial activity of *Pseudonocardia sp. JB05*, a rare salty soil actinomycete against *Staphylococcus aureus*. BioMed Res Int, 2014.

Kämpfer P, Glaeser SP, Parkes L, van Keulen G, Dyson P. 2014. The Family Streptomycetaceae *The Prokaryotes*: Springer, 889–1010.

Kembel SW, Jones E, Kline J, Northcutt D, Stenson J, Womack AM, Bohannan BJ, Brown GZ, Green JL. 2012. Architectural design influences the diversity and structure of the built environment microbiome. ISME J, 6:1469.

Kembel SW, Meadow JF, O’Connor TK, Mhuireach G, Northcutt D, Kline J, Moriyama M, Brown GZ, Bohannan BJ, Green JL. 2014. Architectural design drives the biogeography of indoor bacterial communities. PloS One, 9:e87093.

Koch R. 1884. Die aetiologie der tuberkulose, mittheilungen aus dem laiserlichen gesundheitsampte. 2: 1–88.

Lax S, Smith DP, Hampton-Marcell J, Owens SM, Handley KM, Scott NM, Gibbons SM, Larsen P, Shogan BD, Weiss S. 2014. Longitudinal analysis of microbial interaction between humans and the indoor environment. Science, 345:1048–1052.

Leff JW. 2016. mctoolsr: microbial community data analysis tools. R package version 0.1.1.1.

Loeffler F. 1884. Untersuchungen über die bedeutung der mikroorganismen für die entstehung der Diphtherie beim menschen, bei der taube und beim kalbe. Berlin: K Ges-Amt.

Madden AA, Grassetti A, Soriano JN, Starks PT. 2013. Actinomycetes with antimicrobial activity isolated from paper wasp (Hymenoptera: Vespidae: Polistinae) nests. Environ Entomol, 42:703–710.

Martínez-Porchas M, Villalpando-Canchola E, Vargas-Albores F. 2016. Significant loss of sensitivity and specificity in the taxonomic classification occurs when short 16S rRNA gene sequences are used. Heliyon, 2:e00170.

McCleery RA, Lopez RR, Silvy NJ, Grant WE. 2005. Effectiveness of supplemental stockings for the endangered Key Largo woodrat. Biol Conserv, 124:27–33.

Meadow JF, Altrichter AE, Kembel SW, Kline J, Mhuireach G, Moriyama M, Northcutt D, O’Connor TK, Womack AM, Brown GZ. 2014. Indoor airborne bacterial communities are influenced by ventilation, occupancy, and outdoor air source. Indoor Air, 24:41–48.

Miletto M, Lindow SE. 2015. Relative and contextual contribution of different sources to the composition and abundance of indoor air bacteria in residences. Microbiome, 3:61.

Nguyen NH, Smith D, Peay K, Kennedy P. 2015. Parsing ecological signal from noise in next generation amplicon sequencing. New Phytol, 205:1389–1393.

Ogle DH. 2018. FSA: fisheries stock analysis. R package version 0.8.20.9000.

Oksanen J, Blanchet FG, Kindt R, Legendre P, Minchin PR, O’hara RB, Simpson GL, Solymos P, Stevens MHH, Wagner H. 2013. Package ‘vegan’. Community ecology package. R package version 2.

Pakarinen J, Hyvärinen A, Salkinoja◻Salonen M, Laitinen S, Nevalainen A, Mäkelä MJ, Haahtela T, Von Hertzen L. 2008. Predominance of Gram◻positive bacteria in house dust in the low◻allergy risk Russian Karelia. Environ Microbiol, 10:3317–3325.

Pike C. 1951. Corynebacterial endocarditis: With report of a case due to toxigenic *Corynebacterium diphtheriæ*. J Pathol Bacteriol, 63:577–585.

Platas G, Morón R, González I, Collado J, Genilloud O, Peláez F, Diez MT. 1998. Production of antibacterial activities by members of the family Pseudonocardiaceae: influence of nutrients. World J Microbiol Biotechnol, 14:521–527.

Pohlert T, Pohlert MT. 2018. Package ‘PMCMR’. R package version 1.4.0.

Rainey DG. 1956. Eastern woodrat, Neotoma floridana: life history and ecology. Lawrence, KS: University of Kansas Publications, Museum of Natural History.

Razzauti M, Galan M, Bernard M, Maman S, Klopp C, Charbonnel N, Vayssier-Taussat M, Eloit M, Cosson J. 2015. A comparison between transcriptome sequencing and 16S metagenomics for detection of bacterial pathogens in wildlife. PLoS Negl Trop Dis, 9:e0003929.

Ruiz-Calderon JF, Cavallin H, Song SJ, Novoselac A, Pericchi LR, Hernandez JN, Rios R, Branch OH, Pereira H, Paulino LC. 2016. Walls talk: microbial biogeography of homes spanning urbanization. Science advances, 2:e1501061.

Ruokolainen L, Von Hertzen L, Fyhrquist N, Laatikainen T, Lehtomäki J, Auvinen P, Karvonen AM, Hyvärinen A, Tillmann V, Niemelä O. 2015. Green areas around homes reduce atopic sensitization in children. Allergy, 70:195–202.

Ruokolainen L, Lehtimäki J, Karkman A, Haahtela T, Hertzen Lv, Fyhrquist N. 2017. Holistic view on health: two protective layers of biodiversity. Ann Zool Fennici, 54: 39–49.

Sakoulas G, Moise-Broder PA, Schentag J, Forrest A, Moellering RC, Eliopoulos GM. 2004. Relationship of MIC and bactericidal activity to efficacy of vancomycin for treatment of methicillin-resistant *Staphylococcus aureus* bacteremia. J Clin Microbiol, 42:2398–2402.

Savage AM, Youngsteadt E, Ernst AF, Powers SA, Dunn RR, Frank SD. 2018. Homogenizing an urban habitat mosaic: arthropod diversity declines in New York City parks after Super Storm Sandy. Ecol Appl, 28:225–236.

Steppan SJ, Schenk JJ. 2017. Muroid rodent phylogenetics: 900-species tree reveals increasing diversification rates. PloS One, 12:e0183070.

Team RC. 2015. R: A language and environment for statistical computing [Internet]. Vienna, Austria: R Foundation for Statistical Computing; 2015.

Thoemmes MS, Emerson JB, Leach J, Hedimbi M, Menninger HL, Dunn RR. *In Prep*. The Himba home: insights into global household ecology from the homes of pastoralists.

Thoemmes MS, Stewart FA, Hernandez-Aguilar RA, Bertone MA, Baltzegar DA, Borski RJ, Cohen N, Coyle KP, Piel AK, Dunn RR. 2018. Ecology of sleeping: the microbial and arthropod associates of chimpanzee beds. Royal Soc Open Sci, 5:180382.

US Department of the Interior. 1984. Endangered and threatened wildlife and plants; determination of endangered status for the Key Largo woodrat and the Key Largo cotton mouse. Federal register, 49:171.

US Fish and Wildlife Service. 1999. Multi-species recovery plan for the threatened and endangered species of South Florida. US Fish and Wildlife Service: Vero Beach, FL.

Wang Q, Garrity GM, Tiedje JM, Cole JR. 2007. Naive Bayesian classifier for rapid assignment of rRNA sequences into the new bacterial taxonomy. Appl Environ Microbiol, 73:5261–5267.

